# *scruff*: An R/Bioconductor package for preprocessing single-cell RNA-sequencing data

**DOI:** 10.1101/522037

**Authors:** Zhe Wang, Junming Hu, Evan W. Johnson, Joshua D. Campbell

## Abstract

**Background:** Single-cell RNA sequencing (scRNA-seq) enables the high-throughput quantification of transcriptional profiles in single cells. In contrast to bulk RNA-seq, additional preprocessing steps such as cell barcode identification or unique molecular identifier (UMI) deconvolution are necessary for preprocessing of data from single cell protocols. R packages that can easily preprocess data and rapidly visualize quality metrics and read alignments for individual cells across multiple samples or runs are still lacking.

**Results:** Here we present *scruff*, an R/Bioconductor package that preprocesses data generated from the CEL-Seq or CEL-Seq2 protocols and reports comprehensive data quality metrics and visualizations. *scruff* demultiplexes, aligns, and counts the reads mapped to genome features with deduplication of unique molecular identifier (UMI) tags. *scruff* also provides novel and extensive functions to visualize both pre- and post-alignment data quality metrics for cells from multiple experiments. Detailed read alignments with corresponding UMI information can be visualized at specific genome coordinates to display differences in isoform usage. The package also supports the visualization of quality metrics for sequence alignment files for multiple experiments generated by Cell Ranger from 10X Genomics. *scruff* is available as a free and open-source R/Bioconductor package.

**Conclusions:** *scruff* streamlines the preprocessing of scRNA-seq data in a few simple R commands. It performs data demultiplexing, alignment, counting, quality report and visualization systematically and comprehensively, ensuring reproducible and reliable analysis of scRNA-seq data.

## Background

Single-cell RNA sequencing (scRNA-seq) technologies can profile the transcriptome of individual cells allowing for greater characterization of cellular heterogeneity in complex biological systems. In the past decade, the number of cells profiled by scRNA-seq technologies within a single experiment has grown from a single blastomere(1) to hundreds of thousands of cells(2). The high throughput is achieved by advanced multiplexing strategies where the mRNA molecules from each cell are barcoded with unique oligonucleotide tags embedded within the reverse transcription primers. After synthesis, the cDNA is subsequently amplified using PCR or *in vitro* transcription. These amplification steps prior to sequencing often introduce bias due to different amplification efficiencies for different molecules(3). To alleviate this problem, random molecular barcodes, usually referred to as unique molecular identifiers (UMIs), are often inserted into RT primers which enable the identification of PCR duplicates(4, 5). After barcoding, cells from multiple plates or droplets are sequenced together and then computationally deconvoluted to obtain counts for each individual cell. Samples within a study are often processed in different plates, batches, or runs but ultimately need to be assessed together.

Available computational tools that preprocesses scRNA-seq data generated from CEL-Seq related protocols have limitations. For example, CEL-Seq pipeline(6), umis(7), and UMI-tools(8) do not report any data quality visualizations. zUMIs and scPipe only report limited QC metrics, have limited plotting capabilities, and do not support parallelization(9, 10). Furthermore, there is no tool that can view detailed read alignments with UMI information on specified genomic coordinates in a single cell. For assessment of 10X Genomics data quality, Cell Ranger does not systematically generate and plot quality control metrics support across multiple experiments(2).

*scruff* performs data preprocessing and reports comprehensive QC metrics and visualizations for data generated by CEL-Seq and CEL-Seq2 protocols. It also supports the visualization of read alignment statistics for BAM files from multiple runs generated by Cell Ranger from 10X protocol. *scruff* package reports detailed metrics on measurements on several different aspects of the data, providing a streamlined assessment of quality control.

### Implementation

*scruff* stands for <u>**S**</u>ingle <u>**C**</u>ell <u>**R**</u>NA-seq <u>**U**</u>MI <u>**F**</u>iltering <u>**F**</u>acilitator and is an R/Bioconductor package(11) that demultiplexes cell barcodes, aligns reads to reference genome, and generate gene-level counts with UMI deduplication from scRNA-seq experiments. The main design aim of *scruff* is the ability to 1) preprocess scRNA-seq sequenced reads and generate gene-level count data for individual cells in parallel and 2) summarize and display of comprehensive data quality metrics. *scruff* supports the preprocessing and data quality visualization of sequenced reads from CEL-Seq(12), CEL-Seq2(6) and SORT-seq(13) protocols. Additionally, *scruff* package complements the report of Cell Ranger pipeline by providing the visualization of read alignment information in BAM files from 10X Genomics for multiple runs simultaneously. All functions and procedures in *scruff* package are implemented using R statistical framework.

## Results and discussion

### Cell barcode demultiplexing

The *scruff* pipeline starts with the demultiplexing of paired-end reads in FASTQ format (**Figure 1**). In the `demultiplex` function, cell barcode and UMI sequences are first trimmed from the reads and UMIs are appended to read headers. Reads are filtered according to the Phred quality scores(14, 15) of their corresponding cell barcode and UMI sequences. Reads with Phred scores lower than a user-defined threshold are removed. In the meantime, cell barcodes are mapped to a user-defined whitelist. Reads with cell barcode mismatches exceeding user-defined threshold are excluded. The remaining reads are stored in cell specific FASTQ files and an annotation table is used to keep track of which experiment the cell is from. *scruff* enables the trimming of read sequences by allowing users to only keep certain number of nucleotides for the reads. This is useful if the sequences at 3’ ends have poor quality scores. Parallelization using the BiocParallel(16) package is implemented to allow for simultaneous demultiplexing of multiple samples.

**Figure 1.**
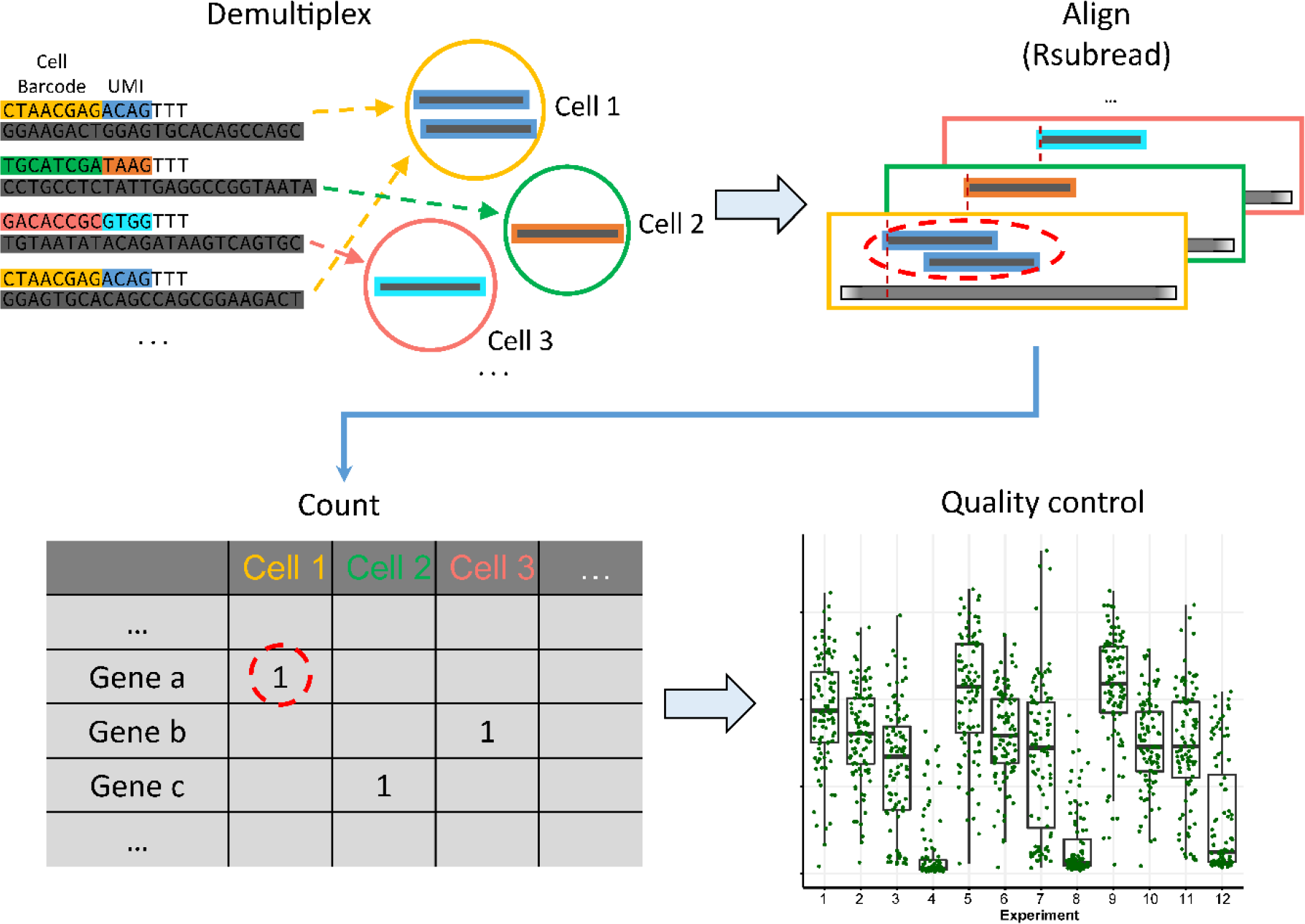
*scruff* package workflow. Reads from FASTQ files are first demultiplexed into cell-specific FASTQ files according to their cell barcodes. During this process, UMI tags are appended to the read header of the transcript sequences. *scruff* applies the Subread(18) algorithm for sequence alignment for each cell. Next, reads mapped to genes are counted according to their UMI. Within each gene, reads with identical UMI (red dashed circle) are counted only once. QC metrics are collected at each of these steps and are used for visualization of data quality.

*scruff* makes use of the `SingleCellExperiment` S4 object(17) as a container for both data and data annotation storage. This object keeps track of the directory to cell-specific demultiplexed FASTQ files and their corresponding cell barcodes. Metrics including the number and fraction of reads assigned to each cell are stored in the cell annotation table of the `SingleCellExperiment` object which is passed to the subsequent alignment step.

### Read alignment

By default, *scruff* package uses the aligner Subread(18) and its corresponding R package Rsubread(19) for read alignment. The function `alignRsubread` is a wrapper function to the align function in Rsubread. It parses the FASTQ file paths from the `SingleCellExperiment` object generated in the demultiplexing step and aligns those files to the reference genome. Sequence alignment files in BAM or SAM formats are generated for each cell and saved to user specified output folder. In the meantime, the locations of these cell specific sequence alignment files and their alignment quality metrics including the number and the fraction of aligned reads are collected and appended to the cell annotation table in the `SingleCellExperiment` object. Reads mapping to multiple genes are removed. Cells can be aligned in parallel to reduce the overall time for this step.

### UMI counting

The `SingleCellExperiment` object containing the file paths to the sequence alignment files are passed to the `countUMI` function for quantification of cell specific gene expression. Genomic features are extracted from the input genome annotation file. *scruff* implements the same counting paradigm as the union counting mode of HTSeq(20). Specifically, read alignments overlapping the exonic regions of exactly one gene are used for transcript counting. After parsing the UMI sequences from the read headers, the number of unique UMIs are summarized for each gene to get the counts for gene-wise mRNA transcripts. External RNA Controls Consortium (ERCC) spike-in RNA controls are flagged by the `isSpike` method so they can be handled separately during data analysis. Gene counting for each cell can be run in parallel. The resulting count matrix is saved in a tab delimited file and stored in the assay slot of the `SingleCellExperiment` object. Gene annotations including gene ID, gene name, and gene biotype are collected from the input gene annotation file and stored in the gene annotation table of the `SingleCellExperiment` object.

In the UMI counting step, various quality metrics are collected for each cell including the number of reads mapped to the genome, number of reads mapped to genes, total number of transcripts (i.e. number of UMIs), number of transcripts from mitochondrial genes, number of transcripts from protein coding genes, number of transcribed genes detected with at least one count, and number of transcribed protein coding genes detected with at least one count. These metrics are appended to the cell annotation table of the `SingleCellExperiment` object. Finally, this `SingleCellExperiment` object containing the count matrix, cell and gene annotation information is returned at the end of the pipeline.

### Modular flexibility

All three preprocessing functions (`demultiplex`, `alignRsubread`, and `countUMI`) are encapsulated in a single function called `scruff` to streamline the entire workflow. The user also has the ability to plug-and-play different methods to generate custom workflow. For example, instead of aligning the reads with Rsubread, users can run other aligners outside of R such as STAR or Bowtie on the demultiplexed FASTQ files. Because the UMI sequences are encoded in the read header in the demultiplexing step, the downstream UMI counting step can be applied to sequence alignment files generated by any alignment algorithms that do not modify the read headers containing the UMI tag.

### Data quality visualization

For scRNA-seq studies, fast and intuitive assessment of read preprocessing quality across experiments and cells is necessary to ensure the validity of downstream data analysis. *scruff* provides functions to visualize both pre- and post-alignment data quality for all cells using quality metrics information collected in demultiplexing, alignment, and UMI counting steps. The `qcplots` function parses the cell annotation table from the `SingleCellExperiment` object and automatically generates numerous boxplots showing various data quality metrics for individual experiments and cells. These metrics include total number of reads, number of reads mapped to reference genome, number of reads mapped to genes, fraction of mapped reads to total reads, fraction of reads mapped to genes to reads mapped to genome, fraction of reads mapped to genes to total number of reads, total number of transcripts, number of mitochondrial transcripts, fraction of mitochondrial transcripts, number of transcribed genes, fraction of protein coding genes, fraction of protein coding transcripts, median and average number of reads per corrected and uncorrected UMI counts, and the number of detected genes divided by total number of reads sequenced per million. These plots can be used for assessing the sample quality of the experiment and across individual batches. From these plots, poor-quality outlier cells can be identified and flagged for removal.

*scruff* package contains method to look at the detailed read alignment information for specific genes in a cell. The function `rview` visualizes all reads mapped at specific region of the chromosome. Function `gview` plots the exonic regions of all gene isoforms between specified genomic coordinates. By combining these two plots vertically using `tracks` function from the ggbio(21) package, users are able to visualize the exact locations of read alignments on the gene.

Finally, *scruff* provides method to visualize read alignment statistics for sequence alignment files from multiple experiments generated by the Cell Ranger pipeline from 10X Genomics. `tenxBamqc` function parses read alignments from sequence alignment files in BAM format. A `SingleCellExperiment` object containing the number of mapped reads and the number of reads mapped to genes for filtered cells is returned. These metrics can be visualized by passing this object to the `qcplots` function.

### Application to example datasets

We tested *scruff* on selected experiments from a publicly available scRNA-seq dataset(22) generated using the CEL-Seq protocol. Raw FASTQ files containing reads from 1,417 single cells across 15 experiments were processed using the `scruff` function. It generates the final `SingleCellExperiment` S4 object containing the transcript count matrix and all cell and gene annotations in one function call. On average, 79,509 reads are sequenced per cell (**Figure 2A**) and 46.78% of reads aligned to GRCm38 (**Figure 2B**). Cells in experiment mouse c library 2 have similar total number of reads compared with other experiments but significantly lower fraction of aligned reads (median is 4.41 %). This is consistent with the mappability (5%) reported in the original study(22). 1,342 cells had greater than 80% nonmitochondrial transcripts (**Figure 2C**) with the median fraction of mitochondrial transcripts at 5.93% indicating that the majority of cells were high quality(23).

**Figure 2.**
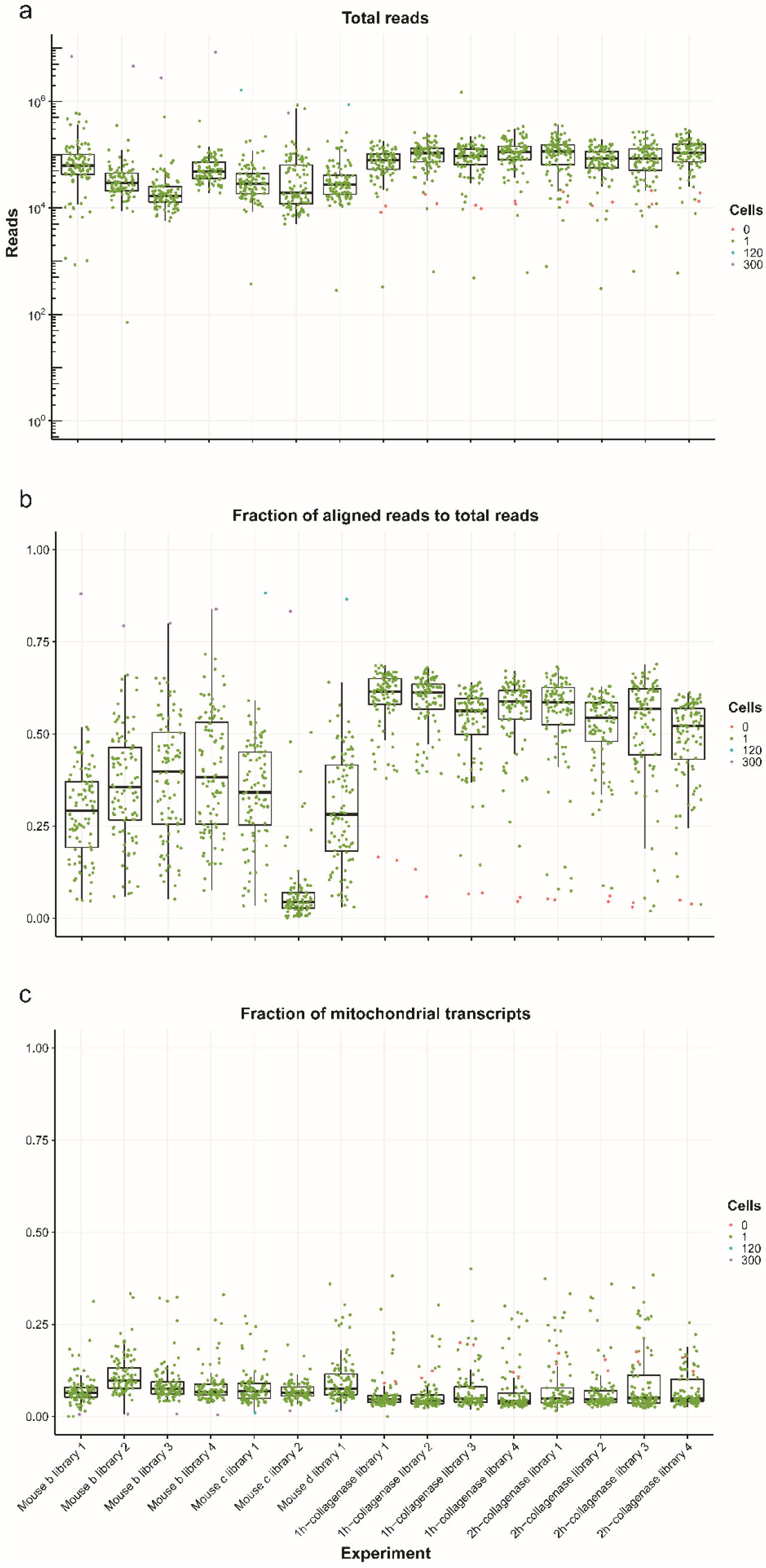
Visualization of example data quality using *scruff*. These are boxplots with overlaying points showing **(a)** the total number of reads, **(b)** the fraction of aligned reads and **(c)** the fraction of mitochondrial transcripts for each experiment. Each point represents a well (unique cell barcode) and is colored by the number of cells sorted in the well by FACS. Each boxplot denotes a different experiment (i.e. plate).

*scruff* package also provides functions to visualize gene isoforms and UMI tagged read alignments at specific genomic coordinates. **Figure 3** shows an example of 125 reads mapped to mouse gene Fos in cell 30 of mouse b library 1. In this case, all of the reads mapped to the 3’ end of transcript Fos-201, indicating the mRNAs transcribed from this gene in this cell are from the isoform Fos-201. The fact that most of the reads are mapped to the 3’ end and forward strand demonstrates that reads sequenced by the CEL-Seq protocol are poly-A selected and maintain strand orientation(12).

**Figure 3.**
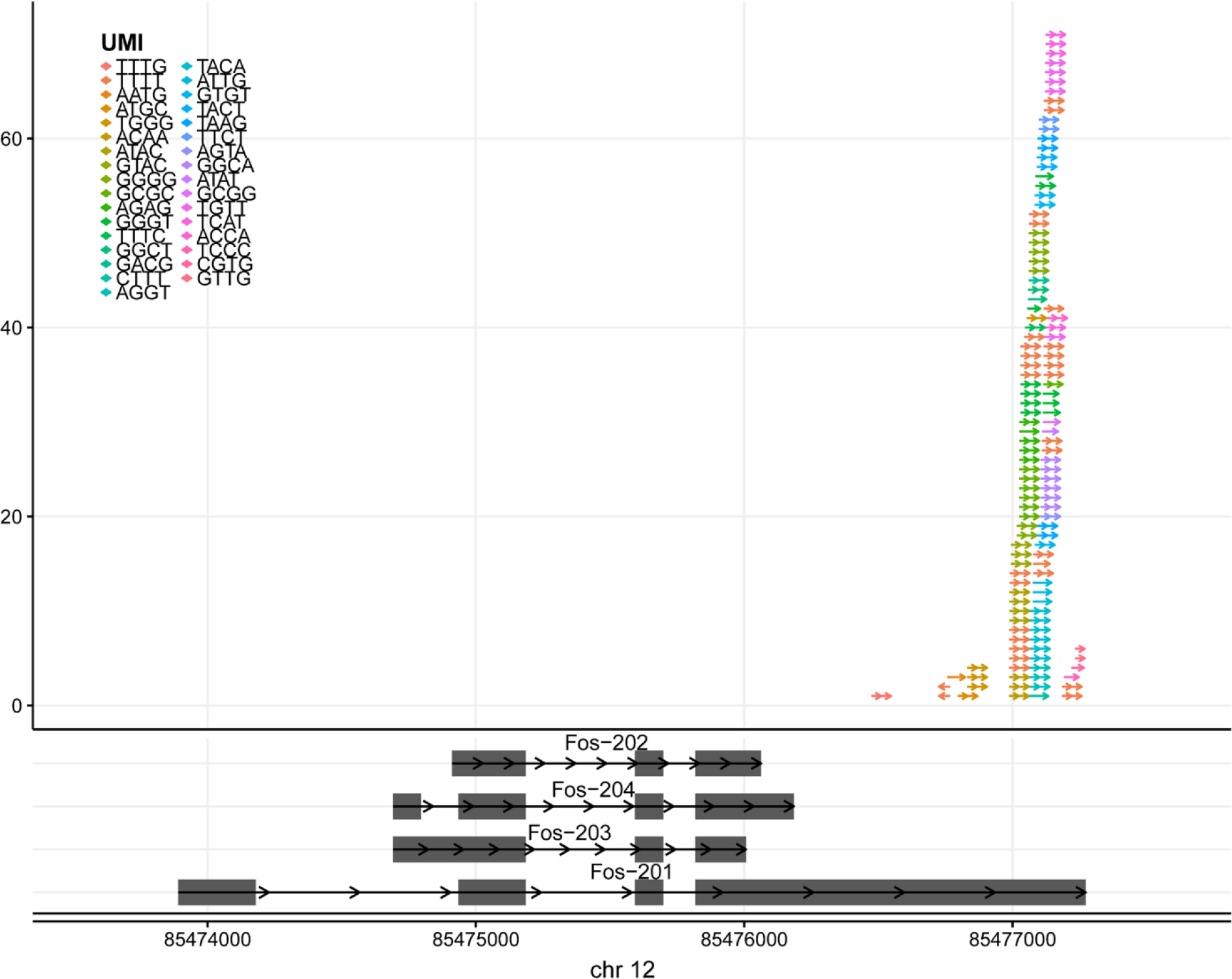
Read alignment visualization with UMI information. 125 reads were mapped to the gene *Fos* in cell 30 of mouse b library 1 from the example CEL-Seq dataset(22). Upper panel shows the visualization of read alignments. Reads are represented by arrows and are colored by their UMIs. The direction of the arrow represents the mapping strand of the read. Lower panel shows the visualization of gene isoforms. Gene isoforms are labeled by their transcript names. Grey rectangles represent exons.

We also applied the read alignment quality visualization function from the *scruff* package to the BAM files of 6 PBMC data downloaded from 10X Genomics website. The number of reads mapped to reference genome, the number of reads mapped to genes, and the fraction of gene reads out of mapped reads were plotted (**Figure 4**). The mean number of aligned reads was 45,000, 92,000, and 56,000 for sequencing libraries prepared with the v1, v2, and v3 reagent kits, respectively. The total number of reads mapped to genes was significantly different between v1 and v2 (p < 2.2 × 10^−16^, two-tailed *t*-test) and v2 and v3 (p < 2.2 × 10^−16^, two-tailed *t*-test) chemistry methods. The proportions of reads mapped to genes were significantly different between v1 and v2 (p = 5.99 × 10^−5^, two-tailed *t*-test), v1 and v3 (p < 2.2 × 10^−16^, two-tailed *t*-test), and v2 and v3 (p < 2.2 × 10^−16^, two-tailed *t*-test) methods. The mean proportion of reads mapped to genes from 1K and 10K PBMC v3 chemistry data is 10.20% lower compared to v1 and v2 chemistry data (p < 2.2 × 10^−16^, two-tailed *t*-test).

**Figure 4.**
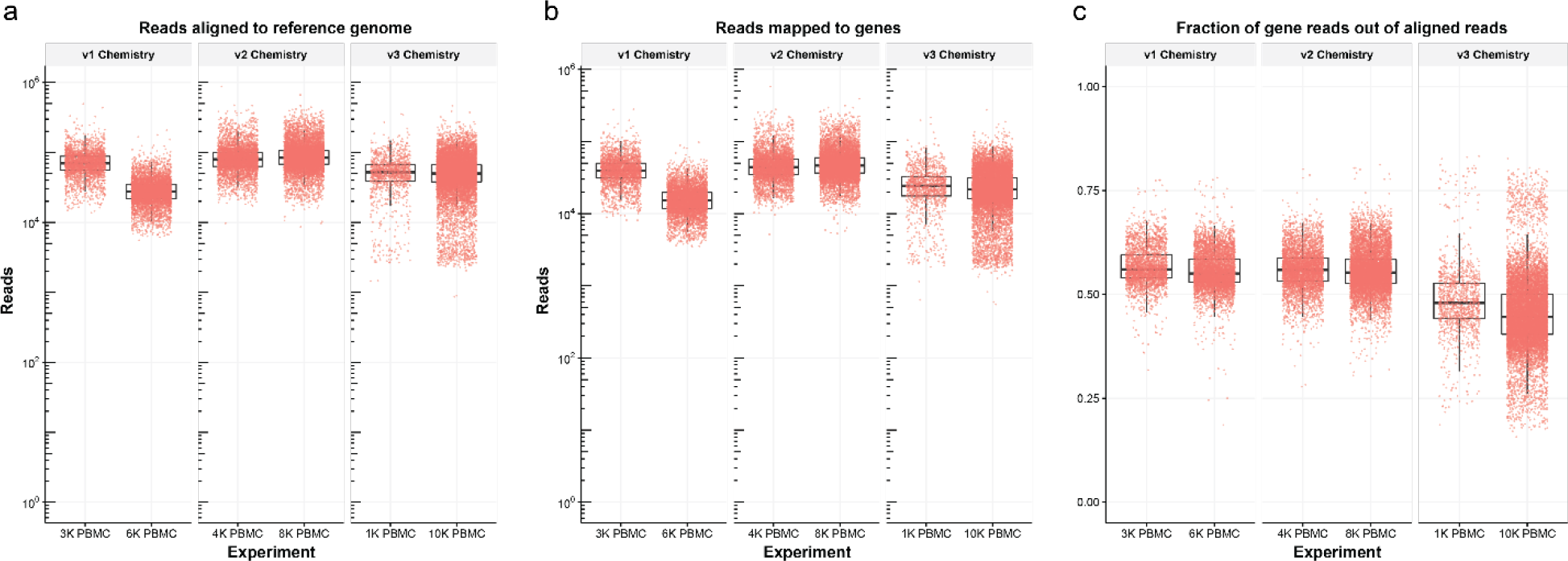
Read alignment quality metrics from 10X Genomics data. BAM files for 6 PBMC datasets were downloaded from 10X Genomics website and processed to obtain **(a)** the number of reads aligned to reference genome, **(b)** the number of reads mapped to genes, and **(c)** the fraction of reads mapped to genes out of total number of aligned reads.

## Conclusions

*scruff* is an R/Bioconductor package that can preprocess scRNA-seq data including demultiplexing cell specific reads, aligning reads to a reference genome, counting the number of transcripts with UMI deduplication, and generating comprehensive plots for quality control across multiple batches or runs. Along with reporting gene expression count matrix as a tab delimited file, *scruff* also promotes data accessibility and portability by generating a `SingleCellExperiment` S4 object which can be passed directly to downstream scRNA-seq data analysis packages including Celda and singleCellTK(24). *scruff* is fully modularized so users can plug and play different tools for performing demultiplexing or alignments. Overall, *scruff* improves single-cell analysis by streamlining preprocessing and quality control workflows.

## Availability and requirements

**Project name**: *scruff*

**Project home page**: http://bioconductor.org/packages/scruff

**Operating system**: Linux, macOS, partially working on Microsoft Windows

**Programming language:** R

**Other requirements**: R >= 3.5, Rsubread.

**License**: MIT License.

## List of abbreviations

UMI: quantitative trait locus
UMI: Unique molecular identifier
scRNA-seq: Single-cell RNA sequencing
PCR: Polymerase chain reaction
SAM: Sequence Alignment Map
BAM: Binary Alignment Map
PBMC: Peripheral Blood Mononuclear Cells
STAR: Spliced Transcripts Alignment to a Reference

## Declarations

### Ethics approval and consent to participate

Not applicable

### Consent for publication

Not applicable

### Availability of data and material

The scruff package is available on Bioconductor (http://bioconductor.org/packages/scruff) and on GitHub (https://github.com/campbio/scruff). The CEL-Seq dataset analyzed in the study are available in the Gene Expression Omnibus repository with accession number GSE85755(22) (https://www.ncbi.nlm.nih.gov/geo/query/acc.cgi?acc=GSE85755). All PBMC BAM files are downloaded from 10X Genomics website (https://support.10xgenomics.com/single-cell-gene-expression/datasets).

## Authors’ contributions

ZW developed the software package; collected, analyzed and interpreted data; and wrote the manuscript. JH assisted in the package’s development, data analysis and visualization. EWJ critically reviewed the manuscript. JDC conceived the project and wrote the manuscript. All authors read and approved the final manuscript.

## Acknowledgements

We thank Prof. Marc Lenburg for helpful suggestions about the development of the package.

